# Hackflex: low cost Illumina Nextera Flex sequencing library construction

**DOI:** 10.1101/779215

**Authors:** Daniela Gaio, Kay Anantanawat, Joyce To, Michael Liu, Leigh Monahan, Aaron E. Darling

**Author notes:** These authors ave contributed equally to this work (Daniela Gaio, Kay Anantanawat). Corresponding author: Daniela Gaio.

## Abstract

We developed a low-cost method for the production of Illumina-compatible sequencing libraries that allows up to 14 times more libraries for high-throughput Illumina sequencing to be generated for the same cost. We call this new method Hackflex. Quality of library preparation was tested by constructing libraries from *E. coli* MG1655 genomic DNA using either Hackflex, standard Nextera Flex or a variation of standard Nextera Flex in which the bead-linked transposase is diluted prior to use. In order to test the library quality for genomes with a higher and a lower GC content, library construction methods were also tested on *P. aeruginosa* PAO1 and *S. aureus* ATCC25923, respectively. We demonstrated that Hackflex can produce high quality libraries and yields a highly uniform coverage, equivalent to the standard Nextera Flex kit. We show that strongly size selected libraries produce sufficient yield and complexity to support *de novo* microbial genome assembly, and that assemblies of the large insert libraries can be much more contiguous than standard libraries without strong size selection. We introduce a new set of sample barcodes that are distinct from standard Illumina barcodes, enabling Hackflex samples to be multiplexed with samples barcoded using standard Illumina kits. Using Hackflex, we were able to achieve a per sample reagent cost for library prep of A$7.22 (USD$5.60), which is 9.87 times lower than the Standard Nextera Flex protocol at advertised retail price. An additional simple modification and further simplification of the protocol by omitting the wash step enables a further price reduction to reach an overall 14-fold cost saving. This method will allow researchers to construct more libraries within a given budget, thereby yielding more data and facilitating research programs where sequencing large numbers of libraries is beneficial.

## INTRODUCTION

The original Nextera protocol provided an easy to use and flexible means for generating Illumina-compatible shotgun libraries. When applied at scale, however, the Nextera reagents at list price could become prohibitively expensive for projects with large sample counts and low sequencing requirements per-sample. Previous work demonstrated that it was possible to dilute the Nextera reagents and with custom buffers, the per-sample library cost could be greatly reduced^1,2^, thereby facilitating the processing of large sample batches. In 2017 Illumina introduced a new type of Nextera kit, called Nextera Flex, and subsequently discontinued the original Nextera kits for which dilution strategies had been developed. The Nextera Flex kits use bead-linked transposases to fragment and tag DNA with adapter sequences. The tagmentation technique allows the incorporation of defined adapter sequences, enabling barcoded primers to anneal and be extended through tagmented DNA fragments in subsequent PCR amplification and sequencing reactions^3^. The new Nextera Flex kits have been shown to yield greatly improved data quality relative to the original Nextera and Nextera XT kits^4^. However, the existing dilution-based protocols^2,3,5^ can not be directly applied with the new Nextera Flex kit, hence we developed an adaptation of those protocols that allows them to work with the Nextera Flex kit protocol, without negative impacts to performance for typical applications such as *de novo* microbial genome assembly.

In this work, we introduce a low cost variant of the Nextera Flex protocol that we call “Hackflex”. In addition to diluting the bead-linked transposases, we propose a simplified protocol that replaces all other reagents with components readily available from third party sources, leading to an 9.87-fold and an 14.17-fold price reduction, with an earlier version of Hackflex (v0) and with the current version of Hackflex (v1), respectively (**Figure 1**; **Supplementary Table 1**). We present our Hackflex protocol and compare the quality of the resulting data to the standard Nextera Flex kit protocol and to a 1:50 bead diluted version of the standard Nextera Flex protocol. We compare the protocols in terms of read quality, uniformity of coverage, coverage of low coverage regions, GC coverage bias, and insert size length. We additionally design 96 barcode sequences that allow the generation of 9120 combinations and we test their performance. Finally, we prepare Hackflex libraries with longer fragment sizes, and demonstrate that large insert Hackflex libraries can lead to improved genome assembly for some microbes.

**Figure 1.**
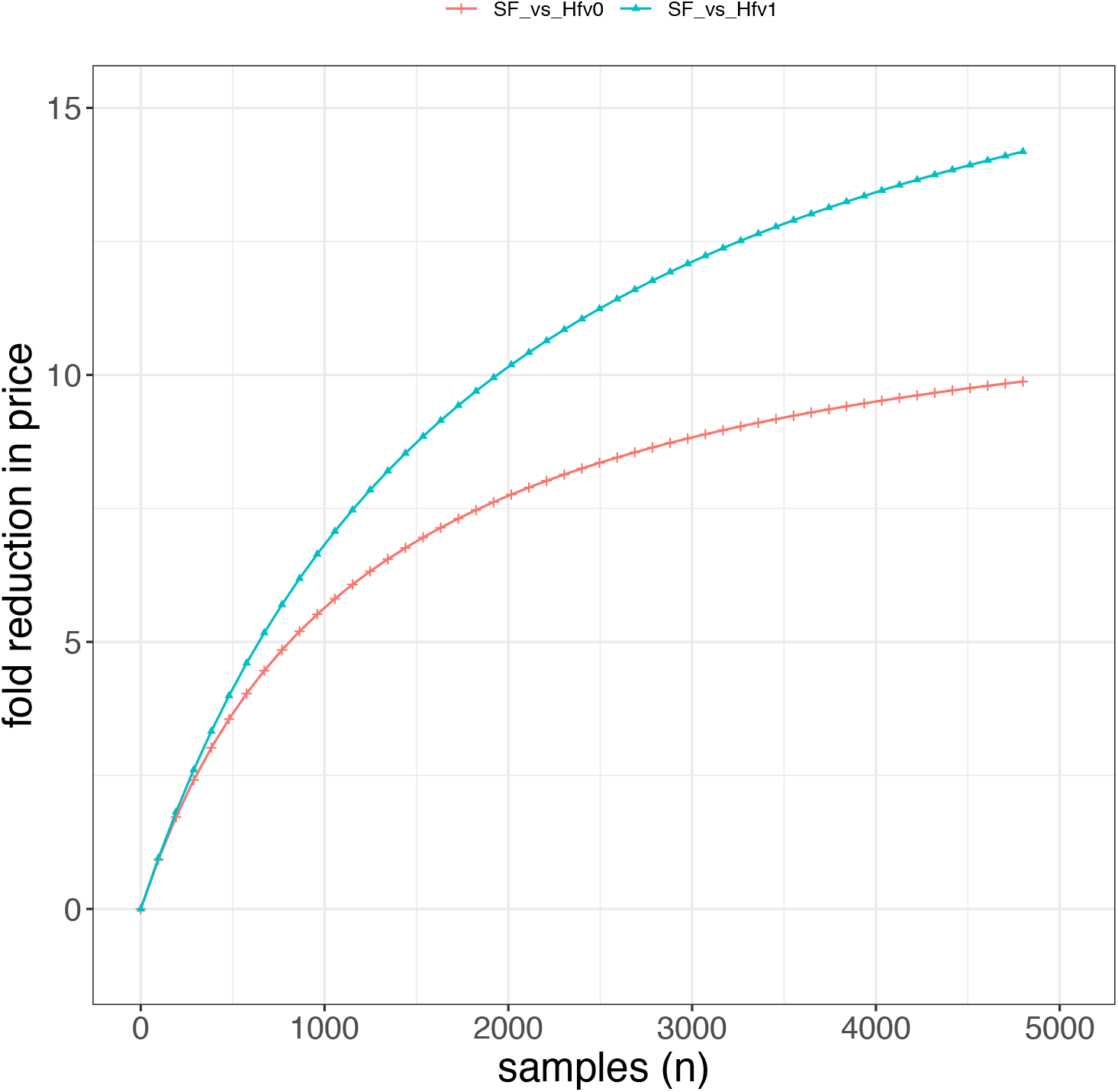
Price fold difference. Fold difference in price between Standard Flex and Hackflex v0 and between Standard Flex and Hackflex v1. Differently from Hackflex v0, Hackflex v1 uses half the amount of the polymerase (1 ul instead of 2 ul) and the TWB wash step is omitted.

## METHODS

### Genomic DNA preparation

Genomic DNA of three different bacteria were used in this study: *Escherichia coli* strain MG1655, *Pseudomonas aeruginosa* strain PAO1, and *Staphylococcus aureus* strain ATCC25923. For *E. coli* MG1655 strain, the reference genome used in this study differs from the original *E. coli* MG1655 strain sequenced by Blattner *et al*^6^, most notably as it contains a pBAD plasmid. Independent reference assemblies of the *E. coli* MG1655 stain, *Pseudomonas aeruginosa* strain PAO1, and *Staphylococcus aureus* strain ATCC25923, were used (see sections “*Nanopore library preparation and sequencing*” and “*Generation of reference assemblies*”). For DNA extraction, high molecular weight gDNA was extracted from freshly cultivated cells of this strain using the Qiagen DNeasy UltraClean Microbial Kit according to the manufacturer’s instructions. Briefly, twenty milliliters of overnight culture was centrifuged at 3200 RCF for 5 min to obtain a cell pellet. Pellets were then washed with 5 mL sterile 0.9% sodium chloride solution, and then resuspended in 300 μL PowerBead solution before continuing with the kit manufacturer’s protocol. Final gDNA was eluted with a 50 μL elution buffer pre-warmed to 42°C. The concentration of isolated DNA samples were measured using a Qubit 2.0 (Thermo Fisher Scientific, USA) and diluted in water.

### Library descriptions

Below we describe the generation of libraries using different protocols and different sources of genomic DNA. The two-letter prefix of each library indicates the source of gDNA used (Ec for *E. coli;* Pa for *P. aeruginosa;* Sa for *S. aureus*). The next two-letter prefix denotes the protocol used (SF: standard Flex; HF: Hackflex). For standard Flex libraries the type of dilution used is described with 1 or 1:50, denoting no dilution (1) or a 1:50 dilution (1:50) (*e.g*.: Ec.SF.B1 *versus* Ec.SF_1:50.B1). Exceptions to the standard protocol are denoted by the letters that follow, up to the suffix. For example, “Ec.SF_PS.B2” indicates that the Standard Flex protocol is used, with PrimeStar GXL polymerase (Takara) as an exception to the standard Flex protocol. The suffix to the library name indicates the source of the sequencing batch (*e.g*.: “Ec.SF.B2” where B2 stands for batch number 2). Each library preparation condition is shown schematically in **Table 1**.

**Table 1.** List of libraries and summary of library preparation conditions.

| Library name | Barcodes | Reagents | Polymerase | Ann. (°C) | Ext. (°C) | Species | BLT dilution | TWB | PCR cycles & Input DNA (ng) | MiSeq cycles |
| --- | --- | --- | --- | --- | --- | --- | --- | --- | --- | --- |
| Ec.SF.B1 | v0(t) | SF | EPM | 62 | 68 | *E. coli* | 1 | NA | 5 - 200 | 600 |
| Ec.SF_1:50.B1 | v0(t) | SF | EPM | 62 | 68 | *E. coli* | 1:50 | NA | 12 -10 | 600 |
| Ec.SF.B2 | v0 | SF | EPM | 62 | 68 | *E. coli* | 1 | NA | 5 - 200 | 600 |
| Ec.SF_1:50.B2 | v0 | SF | EPM | 62 | 68 | *E. coli* | 1:50 | NA | 12 -10 | 600 |
| Ec.SF_PS.B2 | v0 | SF | PS 2ul | 62 | 68 | *E. coli* | 1 | NA | 5 - 200 | 600 |
| Ec.HF.B3 | v0 | HF | PS 1ul | 62 | 68 | *E. coli* | 1:50 | Y | 12 -10 | 600 |
| Ec.HF_55A.B2 | v0 | HF | PS 2ul | 55 | 68 | *E. coli* | 1:50 | Y | 12 -10 | 600 |
| Ec.HF_06x.B3 | v1 | HF | PS 1ul | 62 | 68 | *E. coli* | 1:50_0.6x | N | 12 -10 | 600 |
| Ec.HF-barcode_v0(t).B0 | v0(t) | HF | PS 2ul | 62 | 68 | *E. coli* | 1:50 | Y | 12 -10 | 300 |
| Ec.HF-barcode_v1.B3 | v1 | HF | PS 1ul | 62 | 68 | *E. coli* | 1:50 | N | 12 -10 | 600 |
| Pa.SF.B1 | v0(t) | SF | EPM | 62 | 68 | *P.aeruginosa* | 1 | NA | 5 - 200 | 600 |
| Pa.SF_1:50.B1 | v0(t) | SF | EPM | 62 | 68 | *P.aeruginosa* | 1:50 | NA | 12 -10 | 600 |
| Pa.HF.B2 | v0(t) | HF | PS 2ul | 62 | 68 | *P. aeruginosa* | 1:50 | Y | 12 -10 | 600 |
| Pa.HF_55A.B2 | v0 | HF | PS 2ul | 55 | 68 | *P. aeruginosa* | 1:50 | Y | 12 -10 | 600 |
| Pa.HF_55A72E.B2 | v0 | HF | PS 2ul | 55 | 72 | *P. aeruginosa* | 1:50 | Y | 12 -10 | 600 |
| Pa.HF_06x.B3 | v1 | HF | PS 1ul | 62 | 68 | *P.aeruginosa* | 1:50_0.6x | N | 12 -10 | 600 |
| Sa.SF.B1 | v0(t) | SF | EPM | 62 | 68 | *S. aureus* | 1 | NA | 5 - 200 | 600 |
| Sa.SF_1:50.B1 | v0(t) | SF | EPM | 62 | 68 | *S. aureus* | 1:50 | NA | 12 -10 | 600 |
| Sa.HF.B2 | v0 | HF | PS 2ul | 62 | 68 | *S. aureus* | 1:50 | Y | 12 -10 | 600 |
| Sa.HF_55A.B2 | v0 | HF | PS 2ul | 55 | 68 | *S. aureus* | 1:50 | Y | 12 -10 | 600 |
| Sa.HF_06x.B3 | v1 | HF | PS 1ul | 62 | 68 | *S. aureus* | 1:50_0.6x | N | 12 -10 | 600 |

### Nextera Flex library preparation

We first created a set of libraries using the standard protocol of Nextera Flex (referred to as “Standard Flex” abbrev. “SF”). Each Standard Flex library was constructed using all standard kit reagents from the Nextera DNA Flex Library Prep kit (Illumina, USA), following the manufacturer’s protocol. Briefly, 200 ng input DNA in 30 ul nuclease free water was tagmented by adding 10 ul of Bead Linked Transposase (BLT) and 10 ul of TB1 solution. Each sample was then incubated in the thermocycler at 55 °C for 15 mins, then held at 10°C. After the incubation, 10 ul of TSB solution was added into the tagmentation reaction, and the sample was incubated at 37°C for 15 mins, then held at 10°C. Each sample was then transferred to the magnetic stand to isolate the DNA-BLT complex. The DNA-BLT complex was washed with 100 ul of TWB solution three times. The PCR reaction for library amplification was prepared by mixing 20 ul of Enhanced PCR Mix (EPM) with 20 ul of nuclease free water. The mixture was added into the DNA-BLT complex. 5 ul of each i5 and i7 adapter was added into the PCR reaction. The final volume of the PCR reaction is 50 ul. PCR conditions were set to 68°C for 3 mins, 98°C for 3 mins, followed by 5 cycles of [98°C for 30 sec - 62°C for 30 sec - 68°C for 2 mins], 68°C for 1 mins and held at 10°C. After library amplification, the sample tube was placed onto the magnetic stand. Forty-five ul of the PCR supernatant was mixed with 85 ul of diluted PB reagent (45 ul of PB solution diluted in 40 ul of RSB solution; the final ratio of beads to the PCR supernatant is 0.53X), then incubated at room temperature for 5 mins. The sample tube was then placed on the magnetic stand, and 125 ul of supernatant was transferred into a new sample tube containing 15 ul of undiluted PB (the ratio of the beads to the starting PCR supernatant is 0.7X). The sample was mixed and incubated at room temperature for 5 mins, then the tube was placed on the magnet. The supernatant was discarded, and the beads were washed with 200 ul of fresh 80% ethanol twice. The beads were left to air-dry at room temperature, and were resuspended in 32 ul of RSB solution. The beads were incubated at room temperature for 2 mins. The sample tube was placed on the magnet, and finally 30 ul of eluted library was transferred into a new sample tube. The concentration of eluted library and the library size were measured using Qubit High Sensitivity dsDNA kit (Thermo Fisher Scientific, USA) and the High Sensitivity Bioanalyzer chip (Agilent Technology, USA), respectively. In total, there were 4 standard Flex libraries: two from *E. coli* genomic DNA (referred to as: “Ec.SF.B1” and “Ec.SF.B2”), one from *P. aeruginosa* genomic DNA (“Pa.SF.B2”), and one from *S. aureus* genomic DNA (“Pa.SF.B2”).

The first step in Hackflex development was to test if the BLT beads can be diluted. Diluted standard Flex libraries were generated by following the standard Nextera Flex protocol using the standard reagents (as described above). The quantity of all reagents used in the reaction were unchanged, except for the BLT beads which were diluted 1:50 with nuclease free water prior to use (referred to as “1:50 diluted Flex” abbrev. “SF_1:50”). The amount of input gDNA was adjusted from 200 ng to 10 ng in proportion to the amount of BLT in the reaction, and the number of PCR cycles was increased from 5 to 12 to reflect the amount of DNA-BLT complex in the library amplification step. Of these diluted libraries, we generated in total of four libraries: two libraries from *E. coli* genomic DNA (“Ec.SF_1:50.B1” and “Ec.SF_1:50.B2”), one library from *P. aeruginosa* genomic DNA (“Pa.SF_1:50.B2”), and one library from *S. aureus* genomic DNA (“Sa.SF_1:50.B2”).

As the polymerase used to generate Hackflex libraries is PrimeStar GXL polymerase (Takara), we assessed the impact of the use of a different polymerase in standard Flex libraries, by generating one standard Flex library from *E. coli* genomic DNA, where EPM (Illumina, USA) was replaced with PrimeStar GXL polymerase (Takara, Japan). We refer to this library as “Ec.SF_PS.B2”.

All Standard Flex and modified Standard Flex libraries described above were purified, pooled in equal molarity, diluted to 4 nM and QC’d on the Bioanalyzer (Agilent Technologies, USA). The final pool was sequenced on Illumina MiSeq platform 2×300 bp using the MiSeq Reagent Kit V3 (600 cycles PE) cartridge (Illumina, USA).

### Hackflex library preparation

Hackflex libraries were prepared using laboratory-made and adapted reagents from the Nextera DNA Flex Library Prep kit (Illumina, USA) (**Supplementary Table 1**). All incubation temperatures and times used in the Hackflex protocol were the same as in the Standard Flex protocol except for the PCR amplification step which is optimised for the polymerase.

Briefly, in the tagmentation reaction, BLT beads were diluted 1:50 with nuclease free water (Invitrogen, USA). Ten nanograms of input gDNA in 10 ul ultrapure water (Invitrogen, USA) was mixed with 10 ul of 1:50 diluted BLT, and 25 ul of 2x laboratory-made tagmentation buffer (20 mM Tris (pH 7.6) (Chem-Supply, Australia), 20 mM MgCl (Sigma-Aldrich, Australia), and 50% (v/v) Dimethylformamide (DMF) (Sigma-Aldrich, Australia)). The final volume of the tagmentation reactions was 45 ul. The sample was incubated at 55°C for 15 mins, then held at 10°C. To stop the tagmentation, 10 ul of 0.2% of sodium dodecyl sulphate (SDS; Sigma-Aldrich, Australia) was added into the sample, instead of using TSB. The sample was then incubated at 37°C for 15 mins and then held at 10°C.

Once the tagmentation was completed, the beads were washed three times using 100 ul of washing solution (0.22 μm MF-Millipore™ membrane filtered solution of 10% polyethylene glycol (PEG) 8000 (Sigma-Aldrich, Australia), 0.25M NaCl (Chem-Supply, Australia) in Tris-EDTA buffer (TE) (Sigma-Aldrich, Australia)), instead of TWB. In the later adaptation of the Hackflex protocol (v1), this wash step was found to be unnecessary and therefore was removed.

For library amplification, EPM master mix was replaced with the PrimeSTAR GXL DNA Polymerase kit (Takara, Japan), following the manufacturer’s protocol. Each PCR reaction contains 10 ul of 5x GXL buffer, 4 ul of 10 mM dNTPs, 2 ul of PrimeStar GXL polymerase, and 19 ul of nuclease free water. In a later adaptation of the Hackflex protocol (v1), the amount of PrimeSTAR GXL polymerase used in the reaction was reduced to 1 ul, and the amount of nuclease free water was increased to 20 ul. The PCR mix was added into washed BLT beads. Barcodes were designed to be used for Hackflex libraries (described below). Five microliters of each 10 uM custom synthesized 96-well plate Illumina Adapter Oligos i5 and i7 were added to a final concentration of 1 uM to each reaction. The final volume for the PCR reaction was 45 ul. Library amplification was performed with different conditions from the manufacturer’s recommended protocol, as follows: 3 min at 68°C, 3 min at 98°C, 12 cycles of [45 sec at 98°C – 30 sec at 62°C – 2 min at 68°C], 1 min at 68°C and hold at 10°C. After library amplification, the samples were individually cleaned up. Size selection and purification of the library followed, replacing reagents SPB and RSB with equal volumes of SPRIselect beads (Beckman Coulter, USA) and ultrapure water (Invitrogen, USA) respectively. The concentration of the library was measured with the Qubit HS dsDNA kit (Thermo Fisher Scientific, USA). Fragment size distribution was assessed using the High Sensitivity DNA kit on the Bioanalyzer (Agilent Technologies, USA). Three libraries were created using the Hackflex protocol described above. These libraries were Ec.HF.B3, Pa.HF.B2 and Sa.HF.B2.

We have created two sets of 96 libraries to test the performances of barcodes v0 and v1 (described in “Barcode design” below). The libraries are Ec-HF-barcode_v0(t).B2 and Ec-HF-barcode_v1.B3 respectively. Briefly, 96 libraries were created using Hackflex described above from *E. coli* genomic DNA. After library amplification, 3 ul of each library was pooled into one tube. Then, the entire pool was cleaned up using SPRIselect beads following the protocol described above. The quality of the pooled library was assessed using Qubit HS dsDNA and the Bioanalyzer. Please refer to the section below “Barcode design” for a description of the barcode designs. For sequencing, the final library was diluted and denatured following manufacturer’s instructions, then 4 pM of the pooled library with 5% PhiX v3 control (Illumina, USA) was loaded onto an Illumina MiSeq instrument and sequenced using MiSeq V3 chemistry, generating 2 x 300 bp paired-end reads.

### Hackflex v1 protocol changes

As mentioned above, the Hackflex protocol v1 includes two differences from the Hackflex protocol v0 described above. These differences consist of: 1) the elimination of the TWB washing step, and 2) the reduction of the amount of PrimeStar GXL DNA Polymerase kit (Takara, Japan) from 2 ul to 1 ul per reaction.

### Additional Hackflex libraries

Different polymerases have their own optimised annealing and extension temperatures. In order to fine-tune the annealing and extension temperatures in the Hackflex protocol, we generated three libraries using an annealing temperature of 55°C, instead of 62°C (“Ec.HF_55A.B2”, “Sa.HF_55A.B2”,”Pa.HF_55A.B2”) and one library from *P. aeruginosa* genomic DNA, where both annealing and extension temperatures were modified (55°C annealing and 72°C extension instead of 68°C; library referred to as: “Pa.HF_55A72E.B2”).

An additional three libraries were prepared following the Hackflex protocol with the aim of producing libraries with longer fragments. For this purpose, all steps of the Hackflex protocols were followed as previously described, except for the library purification clean-up step, where the size selection of the library was done using the ratio of beads to the PCR product of 0.6X. Briefly, after library amplification, 45 ul of the library was mixed with a 27 ul solution of SPRI beads (0.6X), then incubated at room temperature for 5 mins. The microtube containing the library was then placed onto a magnetic stand to pellet the beads. The supernatant was discarded, and the beads were washed twice with 200 ul freshly prepared 80% ethanol. The beads were left to air-dry, then resuspended in 30 ul of nuclease free water. The microtube was removed from the magnetic stand, and incubated at room temperature for another 5 mins. The microtube was then placed back onto the magnetic stand to pellet the beads. The purified library was transferred into a new microtube, and brought through the 0.6X size selection step one more time. We refer to these libraries as: “Ec.HF_06x.B3”; “Pa.HF_06x.B3”; “Sa.HF_06x.B3”.

All libraries described here were sequenced on a Miseq V3 600 cycle cartridge, generating 2×300 bp reads.

### Barcode design

With the purpose of up-scaling the number of combinations for Illumina library preparation from 384 libraries to 9,216, we designed barcodes. We generated two designs of barcodes, which we refer to as barcodes v0 (**Supplementary Table 2**) and barcodes v1 (**Supplementary Table 2**). Both designs consist of i5 (n=96) and i7 (n=96) oligo sequences, where each sequence measures 8 bp in length. In the barcode v0 design, the i5 and i7 oligos are the reverse complement sequences of each other, whereas the barcode v1 design does not include complement sequences. As it is not recommended to use the tandem complement barcodes on the Novaseq platform (Illumina tech support, pers. comm.), only 9,120 of the 9,216 possible barcode combinations can be used out of the barcode v0 design barcodes.

Hairpin, homodimer, and heterodimer formation of the entire barcode sequences (F5 + i5 oligo + N5 or F7 + i7 oligo + N7) for both barcode designs (v0 and v1) were examined using the OligoAnalyzer tool from Integrated DNA Technologies (IDT) website interface (https://sg.idtdna.com/pages/tools/oligoanalyzer).

The performance of both barcode designs (v0 and v1) were tested by constructing two sets of 96 Hackflex libraries, where all libraries were generated from *E. coli* genomic DNA, and each library was constructed using a distinct set of barcodes. For both designs, barcode sequences were designed such that no barcode contained 3 or more identical bases in a row. The barcode v0 set of libraries was sequenced on Miseq V3 300 cycles cartridge, 2×150 bp. The barcode v1 set of libraries was sequenced on Miseq V3 600 cycles cartridge, 2×300 bp.

### Hackflex v1 barcode oligo design

Hackflex v1 barcode oligos were designed to satisfy a particular set of design criteria, namely:

➢ Distinct from currently used length 8 barcodes from common library kits (Illumina TruSeq, NEB)
➢ Do not contain homopolymers and certain sequences thought to be associated with sequencing error on some versions of Illumina chemistry^7^ (AAA,CCC,GGG,TTT,ACA,CAC,GTG,TGT,GGCAG,GGCCG,GGCGG,GGCT G)
➢ Have a minimum edit distance of 3 from each other
➢ Do not result in an oligo that forms a hairpin with a Tm > 51C
➢ Do not result in an oligo that forms a homodimer with a Tm > 58C
➢ Do not produce heterodimers between i5 and i7 that exceed a particular threshold Tm

A custom python script was implemented, using the Primer3 python API for thermodynamic calculations^8^, which applied these design criteria to select a final set of 96 x i5 and 96 x i7 Hackflex v1 barcode oligos. The script is available at https://github.com/GaioTransposon/Hackflex.

### Nanopore library preparation and sequencing

The genomic reference of *E. coli* MG1655, against which libraries were mapped, was generated using both Illumina and Oxford Nanopore sequencing data. For long-read MinION sequencing, libraries were prepared using the 1D ligation sequencing kit (SQK-LSK108) from Oxford Nanopore Technologies (ONT) with modifications to the standard ONT protocol as described previously^6^. Samples were barcoded using the Native Barcoding Expansion kit (EXP-NBD103) and barcoded templates were then pooled together with two other samples from an unrelated project. The final library was loaded onto a ONT MinION instrument with a FLO-MIN106 (R9.4) flow cell and run for 48 h as per the manufacturer’s instructions.

### Data availability

All sequence data has been deposited in NCBI under accession number PRJNA549801. The workflow including the library processing steps and the data analysis are schematically illustrated in **Figure 2**.

**Figure 2.**
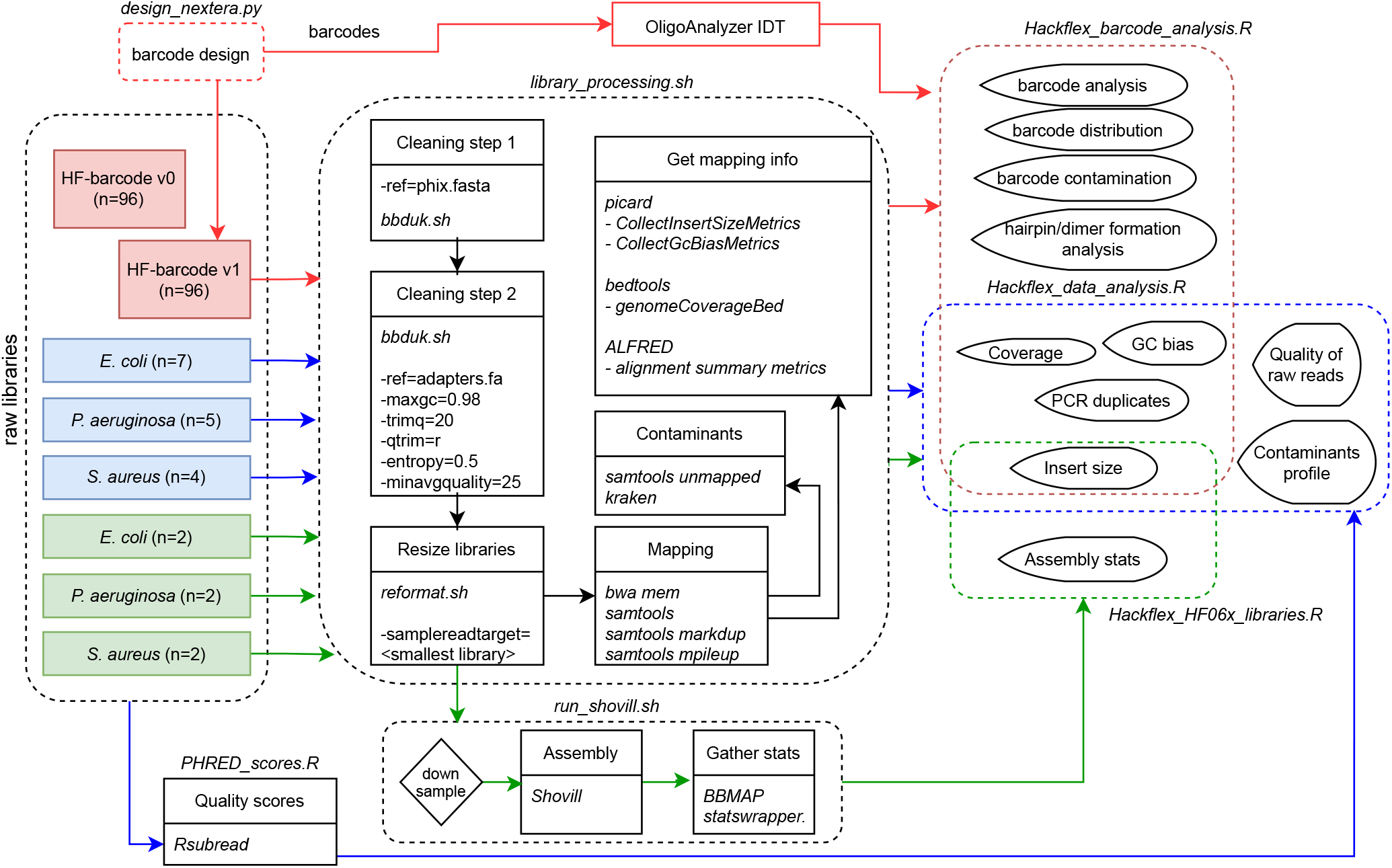
Workflow. Schematic overview of library processing and data analysis performed in this study.

### Barcode demultiplexing

Reads were demultiplexed with bcl2fastq (bcl2Fastq 2.18.0.12, Illumina, Inc.) software with default settings, allowing one mismatch per index. Barcode counts were retrieved from the demultiplexing statistics output of bcl2fastq and histograms representing barcode distribution were generated with R Studio, version 1.1.463 (RStudio: Integrated Development Environment for R, Boston, Massachusetts). Barcode cross-contamination was assessed for barcodes v1 libraries. Barcode cross-contamination was determined by demultiplexing the data, using all possible barcode v1 combinations (n=9216). The rate of barcode cross-contamination is determined as the number of reads attributed to unexpected combinations (i5-i7 combos not used for the run) over the total number of reads assigned to both expected and unexpected combinations. In this manner, it is possible to distinguish the proportion of reads that failed to demultiplex because of sequencing error from those that contributed to barcode cross-contamination.

### Processing of libraries before mapping

Raw reads were assessed for quality with FastQC version 0.11.8 (http://www.bioinformatics.babraham.ac.uk/projects/fastqc/) and using the Bioconductor package Rsubread^6^. Libraries were processed using the bbtools package (http://jgi.doe.gov/data-and-tools/bbtools) (version 38.22). PhiX removal was performed (parameters: `k=31 hdist=1`) and adapter removal, quality trimming and filtering, were performed on all the libraries with BBDuk (parameters: `ktrim=r k=21 mink=11 hdist=1 tpe tbo maxgc=0.98 qtrim=rl qtrim=20 entropy=0.5`). Libraries were downsampled with the bbmap script *reformat.sh* to match the total number of bases of the smallest library included in the comparison group (parameter - `samplebasestarget`).

### Generation of reference assemblies

Reference assemblies for library QC were generated either via de novo assembly of short reads or via hybrid assembly with Nanopore sequence data, where available. For *E. coli*, nanopore data was coassembled with Nextera Flex libraries Ec.SF.B1 and Ec.SF_1:50.B1 with the Unicycler software (version 0.46) with default parameters in “normal” mode. For *P. aeruginosa*, the Nextera Flex libraries Pa.SF.B1 and Pa.SF_1:50.B1 were used. The final reference genomes obtained from the assemblies consisted of 5 (L50=1; N50=4,640,167), and 47 (L50=6; N50=350,586) contigs for *E. coli*, and *P. aeruginosa*, respectively. For *S. aureus*, the reference genome available at SAMN03009478 was used, which consisted of 2 contigs (L50=1; N50=2,778,854).

### Assembly of longer insert size Hackflex libraries

The three double-left clean up Hackflex libraries (Ec.HF_06x.B3, Pa.HF_06x.B3, Sa.HF_06x.B3), together with the Hackflex libraries (Ec.HF.B3, Pa.HF.B2, Sa.HF.B) were resized with *reformat.sh* (as described above), as to obtain an equal number of base pairs for each library, and they were separately *de novo* assembled using shovill^9^ (version 1.1.0). Assembly was performed with libraries downsampled at several depths (parameter `--depth`): 5x, 10x, 20x, 50x, 100x, 150x, 200x. Stats were run on the produced assemblies with *statswrapper.sh* (http://jgi.doe.gov/data-and-tools/bbtools) (version 38.22).

### Short read mapping and coverage analysis

In order to assess performance of the library prep methods, libraries were aligned with bwa mem^10^ (http://jgi.doe.gov/data-and-tools/bbtools) (version 38.22) alignment software and samtools^11^ (version 1.9) (http://samtools.sourceforge.net/) to the reference genomes. After mapping, PCR duplicates were removed using the samtools command `markdup`. Samtools `mpileup` was used to generate the mapped depth of coverage information for each position on the reference genomes. (**Figure 2**)

Insertion fragment data was obtained with picard.jar *CollectInsertSizeMetrics* (version 2.25.0-5) (http://broadinstitute.github.io/picard/) (parameters: M=0.4). Coverage by GC content was obtained with picard.jar *CollectGcBiasMetrics* (version 2.25.0-5) (http://broadinstitute.github.io/picard/) (default parameters). Coverage across the genome was obtained with picard.jar *CollectWgsMetrics* (version 2.25.0-5) (http://broadinstitute.github.io/picard/) (parameters: `READ_LENGTH=300`). Bedgraphs to report coverage across all sites of the reference genomes were obtained with genomeCoverageBed (bedtools v2.27.1) (parameters: `-d -ibam`). Mapping files were processed with ALFRED^12^ from which the alignment summary metrics were extracted. Reads that failed mapping were extracted with samtools view (version 1.9) (http://samtools.sourceforge.net/) (parameters: `-f 0×4`) and analysed with Kraken2^13^ (version 2.0.8-beta) against the minikraken2_v1_8GB database (default parameters). (**Figure 2**) The contaminants profiles are reported in **Supplementary Table 2**.

All data was processed and visualized with R Studio, version 1.1.463 (RStudio: Integrated Development Environment for R, Boston, Massachusetts). All scripts used for the analysis and generation of plots are available at https://github.com/GaioTransposon/Hackflex.

## RESULTS

### Quality of raw reads

Quality scores from all libraries were analysed using the Bioconductor package Rsubread^14^. PHRED scores were similar between Standard Flex and Hackflex libraries, for all libraries made from *E. coli, P. aeruginosa*, and *S. aureus* genomic DNA, with the exception of standard Flex and 1:50 Flex *E. coli* libraries where batch 1 libraries had a lower average PHRED score and a higher standard deviation compared to batch 2 libraries (**Supplementary Table 2**).

### How adapter trimming and quality filtering affected distinct libraries

The cleaning step, consisting of adapter trimming and quality filtering, affected libraries made from *E. coli* genomic DNA with either Standard Flex or the Hackflex protocol in a similar manner, where 3.65%, 3.06%, and 3.31% of the reads were removed from Standard Flex, 1:50 diluted Flex, and Hackflex, respectively. A higher percentage of reads were removed from batch 2 than from batch 1 libraries (Ec.SF.B1: 3.65%; Ec.SF_1:50.B1: 3.06%; Ec.SF.B2: 5.42%; Ec.SF_1:50.B2: 4.80%). Libraries from *P. aeruginosa* genomic DNA had 4.64%, 5.26% and 4.67% of the reads removed during the cleaning step, for Standard Flex, 1:50 diluted Flex, and Hackflex respectively. Libraries from *S. aureus* genomic DNA had a higher percentage of reads removed from the Hackflex (3.14%) compared to Standard Flex (1.74%) and 1:50 diluted Flex (1.97%) library. (**Table 2**) In order to compare the performance of the library protocols, within each species all libraries were resized based on the smallest library, as to obtain an equal number of base pairs for each library (**Table 2**). We report the average read length, the number of reads and base pairs obtained from each library, before and after the cleaning steps, in **Supplementary Table 2**.

**Table 2.** Overview of raw libraries, quality scores, number of reads obtained and filtered, with the standard Flex, diluted Flex, and Hackflex.

| Library | Phred score<br>(mean) | Phred score<br>(sd) | Read count | Removed reads<br>(%) | Size after resizing<br>(bp) |
| --- | --- | --- | --- | --- | --- |
| Ec.SF.B1 | 30.47 | 6.01 | 1000824 | 3.65 | 174760709 |
| Ec.SF_1:50.B1 | 29.9 | 6.85 | 1200710 | 3.06 | 175814069 |
| Ec.SF.B2 | 33.24 | 3.86 | 1837948 | 5.42 | 175540961 |
| Ec.SF_1:50.B2 | 32.33 | 4.68 | 1955102 | 4.8 | 175569760 |
| Ec.HF.B3 | 33.1 | 5.38 | 1486078 | 3.31 | 175506039 |
| Pa.SF.B1 | 28.95 | 6.93 | 974714 | 4.64 | 148086604 |
| Pa.SF_1:50.B1 | 28.29 | 7.4 | 1581282 | 5.26 | 149084666 |
| Pa.HF.B2 | 31.87 | 4.89 | 1631262 | 4.67 | 148779715 |
| Sa.SF.B1 | 32.87 | 4.53 | 779660 | 1.74 | 166527991 |
| Sa.SF_1:50.B1 | 32.17 | 5.15 | 1090564 | 1.97 | 167353079 |
| Sa.HF.B2 | 33.95 | 3.46 | 4236794 | 3.14 | 167220961 |

Given the relatively small amount of sequencing data required for microbial isolate genome sequencing, PCR duplicates would be expected to be very rare in standard Nextera Flex reactions, which are capable of preparing human genomes for shotgun sequencing (30x coverage of a 3Gbp genome). However, by reducing the amount of BLT and increasing the number of PCR cycles used in Hackflex the library complexity is expected to be reduced, and therefore a higher incidence of PCR duplicates may occur. To understand the impact of Hackflex on PCR duplicates, we compared the PCR duplicates obtained with Standard Flex, 1:50 diluted Flex, and Hackflex. We do not observe an excess of PCR duplicates in 1:50 diluted and Hacklex libraries compared to standard Flex libraries, at least at the sequencing depth tested (**Table 3**; **Supplementary Table 2**).

**Table 3.** Mapping details for libraries generated with standard Flex, diluted Flex, and Hackflex.

| Library | Unmapped reads | Mapped Fraction | Mismatch Rate | PCR duplicates | Mean Coverage | SD Coverage |
| --- | --- | --- | --- | --- | --- | --- |
| Ec.SF.B1 | 170 | 0.989 | 0.002 | 10906 | 36.22 | 13.41 |
| Ec.SF_1:50.B1 | 164 | 0.992 | 0.002 | 8655 | 36.55 | 12.77 |
| Ec.SF.B2 | 53 | 0.991 | 0.001 | 6998 | 36.4 | 13.08 |
| Ec.SF_1:50.B2 | 134 | 0.993 | 0.001 | 5288 | 36.76 | 11.14 |
| Ec.HF.B3 | 114 | 0.994 | 0.001 | 4738 | 36.46 | 12.49 |
| Pa.SF.B1 | 107 | 0.994 | 0.002 | 5678 | 23.4 | 7.14 |
| Pa.SF_1:50.B1 | 160 | 0.995 | 0.002 | 4335 | 23.55 | 7.22 |
| Pa.HF.B2 | 73 | 0.996 | 0.001 | 2918 | 23.65 | 7.84 |
| Sa.SF.B1 | 48 | 0.988 | 0.002 | 9255 | 58.56 | 14.32 |
| Sa.SF_1:50.B1 | 123 | 0.991 | 0.001 | 7674 | 58.93 | 14.18 |
| Sa.HF.B2 | 237 | 0.993 | 0.001 | 4627 | 59.03 | 15.14 |

### Coverage

Coverage profiles were obtained by mapping the equally re-sized libraries against their reference genomes. As expected, a highly similar median coverage was obtained among libraries made using the different protocols, independently of the genomic DNA source. Mapped read fractions, average coverage depth, and other mapping metrics for *E. coli, P. aeruginosa*, and *S. aureus* libraries were similar among Standard Flex, diluted Flex and Hackflex libraries (**Table 3**; **Supplementary Table 2**).

### Low coverage regions

Standard flex (batch 1 and 2), 1:50 flex (batch 1 and 2), and Hackflex libraries from *E. coli* genomic DNA had no sites with zero coverage. Libraries made from *S. aureus* genomic DNA had 0, 0, and 1 sites with zero coverage, for Standard Flex, 1:50 Flex, and Hackflex libraries, respectively. All sites with zero coverage of the *E. coli* and *S. aureus* libraries were located near the extremities of contigs (**Supplementary Table 2**). Libraries made from *P. aeruginosa* genomic DNA had considerably more sites with zero coverage than libraries from *E. coli* and *S. aureus*, possibly due to the fragmented reference genome. In fact, there were 651, 263, 13,341 sites with zero coverage, for Standard Flex, 1:50 Flex, and Hackflex libraries, respectively. The low coverage regions were overlapping to a certain extent among the three libraries, possibly indicating common features of these regions that bias against their sequencing with Illumina chemistry. Genomic sites with zero coverage for all libraries are reported in **Supplementary Table 2**.

### Barcode distribution and quality

Two sets of ninety-six i5 and i7 oligos (8 bp), referred to as barcodes v0 and v1, were designed for this study to provide a resource for high throughput multiplexing of Hackflex libraries. To this end, performance of the two sets of 96 designed barcodes was evaluated by subjecting *E. coli* MG1655 DNA to 2×96 independent library constructions with Hackflex reagents, each library with a different barcode combination. Barcode distribution for both barcode designs and GC bias is displayed in **Supplementary Figure 1**.

### Barcodes v0

The 96 libraries with the barcode v0 design are made using the tandem complement design, and we refer to these libraries as Ec.HF-barcode_v0(t).B0, where “v0” is to indicate the barcode design (v0) and the “(t)” is to indicate that these libraries were constructed using i5 and i7 oligos that are tandem complement of each other. Of these 96 libraries, 2 had failed (read count n=12 and n=18). Average barcode count was 11,033.8 (sd=4,149.3). Coefficient of variation was 0.38 and 0.34 including all libraries and excluding the two failed libraries, respectively. The average GC content of oligos is 49.87 (median=50) and two of the 192 v0 oligos have a GC content of 87.5%. The GC content of the entire barcode sequence (*i.e*.: F5+i5+N5 or F7+i7+N7), was plotted against the read count obtained from each barcode, to estimate the extent of GC bias. A significant correlation between read counts obtained and GC content was found for i5 (*R*=-0.704; *p*<0.0001) and i7 (*R*=-0.704; *p*<0.0001) (**Supplementary Figure 1**).

Barcodes v0 had an estimated ΔG between −10.7 and −4.2 kcal.mole-1, and their average melting temperature was 50.36°C (sd=4.56), which is less than 10 degrees lower than the annealing temperature used in the PCR reaction (62°C) (**Table 1**). We examined the hairpin formation, homo- and heterodimer formation of the entire barcodes used to build the two failed libraries. These libraries had a normal estimated deltaG (mean=-6.45 sd=0.73) and a melting temperature that approached the 62°C annealing temperature used in the Hackflex protocol (C3_i5=55.5°C; C3_i7=48.6°C; H7_i5=51.2°C; H7_i7=58.1°C). These two libraries were analyzed for hairpin, homodimer and heterodimer formation. A hairpin formation was found for an i5 oligo (C3_i5 ΔG=-5.5; Tm=55.5°C) and for one i7 oligo (H7_i7 ΔG=-7.02; Tm=58.1°C), possibly explaining the failure of these two libraries (**Supplementary Table 2**).

### Barcodes v1

The 96 libraries made from barcodes v1 do not follow the tandem complement design. The average barcode count was 221,203.1 (sd=53,836.6). Coefficient of variation was 0.243 and 0.183 including all libraries and excluding the three libraries with low counts (8,036; 32,469; 43,836), respectively. The average GC content of oligos is 39.65 (median=50) and two v1 oligos have a GC content of zero. The GC content of the entire barcode sequence (*i.e*.: F5+i5+N5 or F7+i7+N7) was plotted against the read count obtained from each barcode, to estimate the extent of GC bias. No correlation was found between read counts obtained and GC content of either i5 (*R*=-0.17; *p*=0.1) or i7 barcode (*R*=0.03; *p*=0.77) (**Figure 3**). The observed barcode cross-contamination rate was 0.002. Assuming uniform cross contamination, this would result in a sample misassignment rate ranging between 0 and 0.2% of reads, depending on which and how many barcodes are used in a pool of samples.

**Figure 3.**
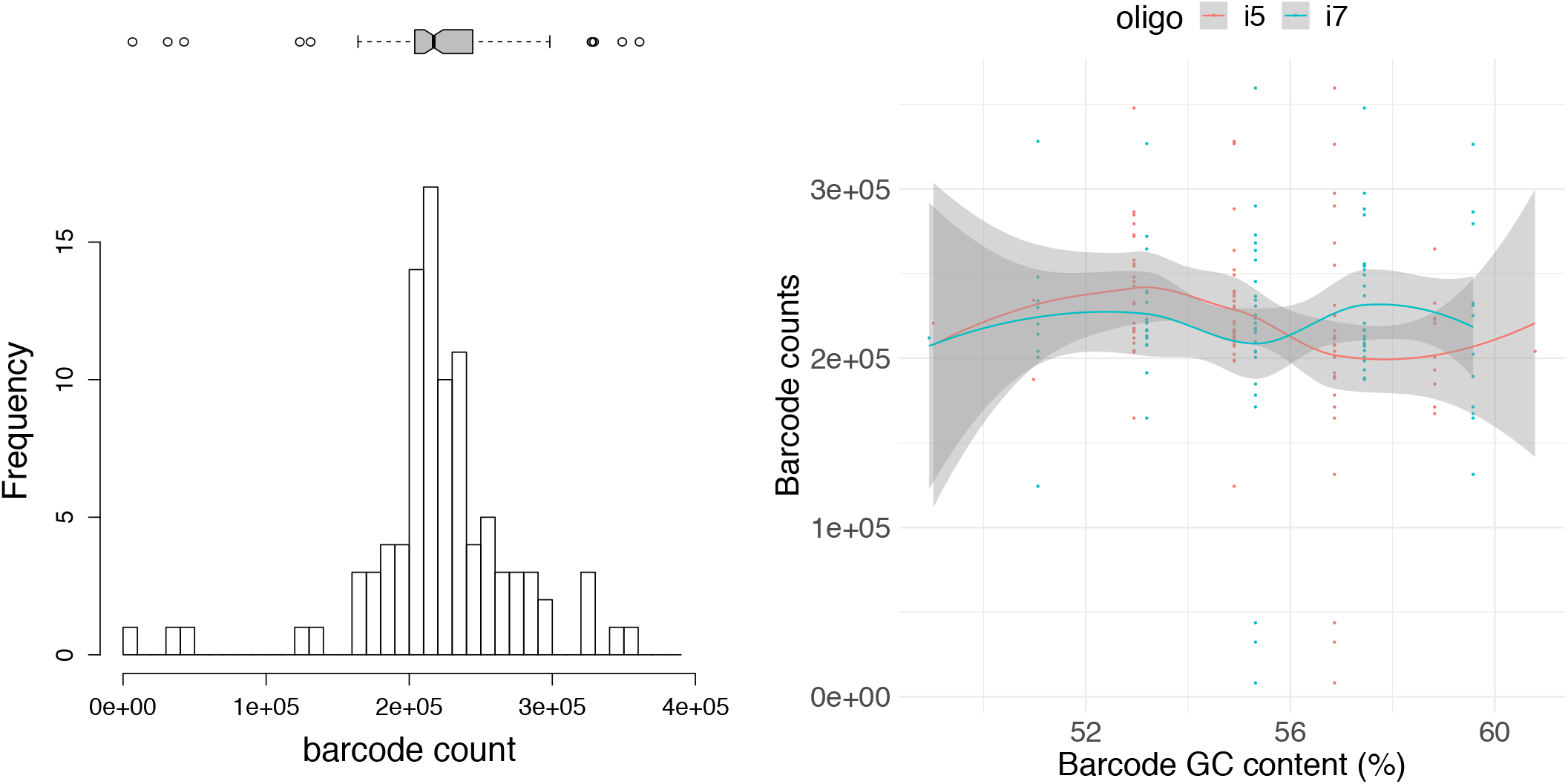
Barcode distribution and GC bias of Hackflex barcode v1 libraries. Unique barcode distribution across 96 Hackflex libraries constructed from barcodes v1(left) and their GC bias: the relation between barcode counts obtained for each entire barcode (i.e.: F5+i5+N5 or F7+i7+N7), and its GC content (i5 *R*=-0.17 p=0.1; i7 *R*=0.03 p=0.77) (right).

The estimated ΔG ranged between −7.660 and −4.220 kcal.mole-1, and the average melting temperature was 49.07°C (sd=3.32), which is over 10 degrees lower than the annealing temperature used in the PCR reaction (62°C) (**Table 1**). We examined the hairpin formation, homo- and heterodimer formation of the entire barcodes used to build the three libraries that yielded a low read count. These libraries had similar ΔG (mean=- 5.72 sd=1.2) and similar melting temperatures (mean=49.6°C sd=1.24) to the rest of the libraries. However, one i5 oligo (v1_A1_i5) was found to form hairpins (ΔG=-7.06; Tm=51.6°C) and one i7 oligo (v1_G2_i7) was found to form a homodimer structure consisting of homodimerization with itself (5 bp) and homodimerization between F7 and N7 (5 bp) in its proximity (ΔG=-13.41; Tm=48.6°C) (**Supplementary Table 2**).

### Insert size

Insert size distribution was analyzed for all libraries. Small differences in insert size distribution were detected between different batches, and although standard Flex libraries from *P. aeruginosa* and *S. aureus* had larger insert sizes compared to diluted 1:50 libraries, an equally large or larger insert size was found in Hackflex libraries compared to standard Flex libraries, in all species (**Table 4**).

**Table 4.** GC content coverage bias and insert size lengths for libraries generated with standard Flex, diluted Flex, and Hackflex.

| Library | Insert size (median) | Insert size (mean) | GC bias (Pearson's R) | *p* value |
| --- | --- | --- | --- | --- |
| Ec.SF.B1 | 302 | 296.31 | -0.886 | *** |
| Ec.SF_1:50.B1 | 230 | 238.68 | -0.8 | *** |
| Ec.SF.B2 | 326 | 334.12 | -0.383 | ** |
| Ec.SF_1:50.B2 | 291 | 296.16 | -0.598 | *** |
| Ec.HF.B3 | 287 | 300.77 | -0.763 | *** |
| Pa.SF.B1 | 269 | 266.97 | -0.965 | *** |
| Pa.SF_1:50.B1 | 241 | 242.39 | -0.965 | *** |
| Pa.HF.B2 | 341 | 351.72 | -0.248 | * |
| Sa.SF.B1 | 293 | 293.1 | 0.698 | *** |
| Sa.SF_1:50.B1 | 269 | 270.79 | 0.099 | ns |
| Sa.HF.B2 | 296 | 304.74 | 0.741 | *** |

The additional ninety-six HF barcode v1 libraries that were generated from *E. coli* DNA to assess the performance of barcodes, underwent a pooled bead clean up and were analyzed to assess the insert size distributions obtained with the Hackflex protocol. With the exclusion of 3 libraries with low counts, 93 libraries had an average insert size of 310.60 base pairs (range of library means: 297.66-314.57; sd range: 120.66-133.29) (**Supplementary Table 2**).

### GC content bias

In order to assess the extent of GC content bias, correlations were obtained between coverage and GC content across the genome using the weighted Pearson’s correlation coefficient (*R*) on the relative (observed/expected) measures of sequence coverage by the reads at each GC content window. All libraries showed to a certain extent a bias at extreme GC content areas as it would be expected^15,16^. The dilution factor and hence number of PCR cycles did not have an obvious association with strength of GC bias, but the GC bias seemed to be associated with batch effects (Ec.SF.B1 *R*=-0.886; Ec.SF_1:50.B1 *R*=-0.8; Ec.SF.B2 *R*=-0.383; Ec.SF_1:50.B2 *R*=-0.598; Pa.SF.B1 *R*=- 0.965; Pa.SF_1:50.B1 *R*=-0.965). In libraries from *S. aureus* genomic DNA, the GC bias was similar between standard Flex and Hackflex (Sa.SF.B1 *R*=0.698; Sa.HF.B2 *R*=0.741; Test statistic z=-0.509 *p*=0.305), and in libraries from *P. aeruginosa* genomic DNA, the GC bias was lower for Hackflex than for standard Flex or 1:50 Flex (Pa.SF.B1 *R*=-0.965; Pa.SF_1:50.B2 *R*=-0.965; Pa.HF.B2 *R*=-0.248; Test statistic z=-10.037 *p*<0.0001) (**Table 4**)

Again, as for the insert size distribution range of Hackflex, we used the HF barcode v1 libraries from *E. coli* genomic DNA to estimate the GC bias with the Hackflex protocol on a large number of libraries. As a high coverage depth can offer a better estimate of GC bias, we included the top twenty largest Hackflex barcode v1 libraries in the analysis (mean depth after resizing of libraries: 12.5x). The average GC bias (Pearson’s *R*) found was −0.642 (sd=0.043; var=0.00188; median=-0.650).

### Size selection on Hackflex libraries can improve de novo assembly

We generated three Hackflex libraries by performing a double left bead clean up. As a result of the double left bead clean up, a median 287-341 insert size length was obtained from Hackflex libraries, and a median 416-433 insert size length was obtained from double-left clean up Hackflex libraries (**Figure 4**).

**Figure 4.**
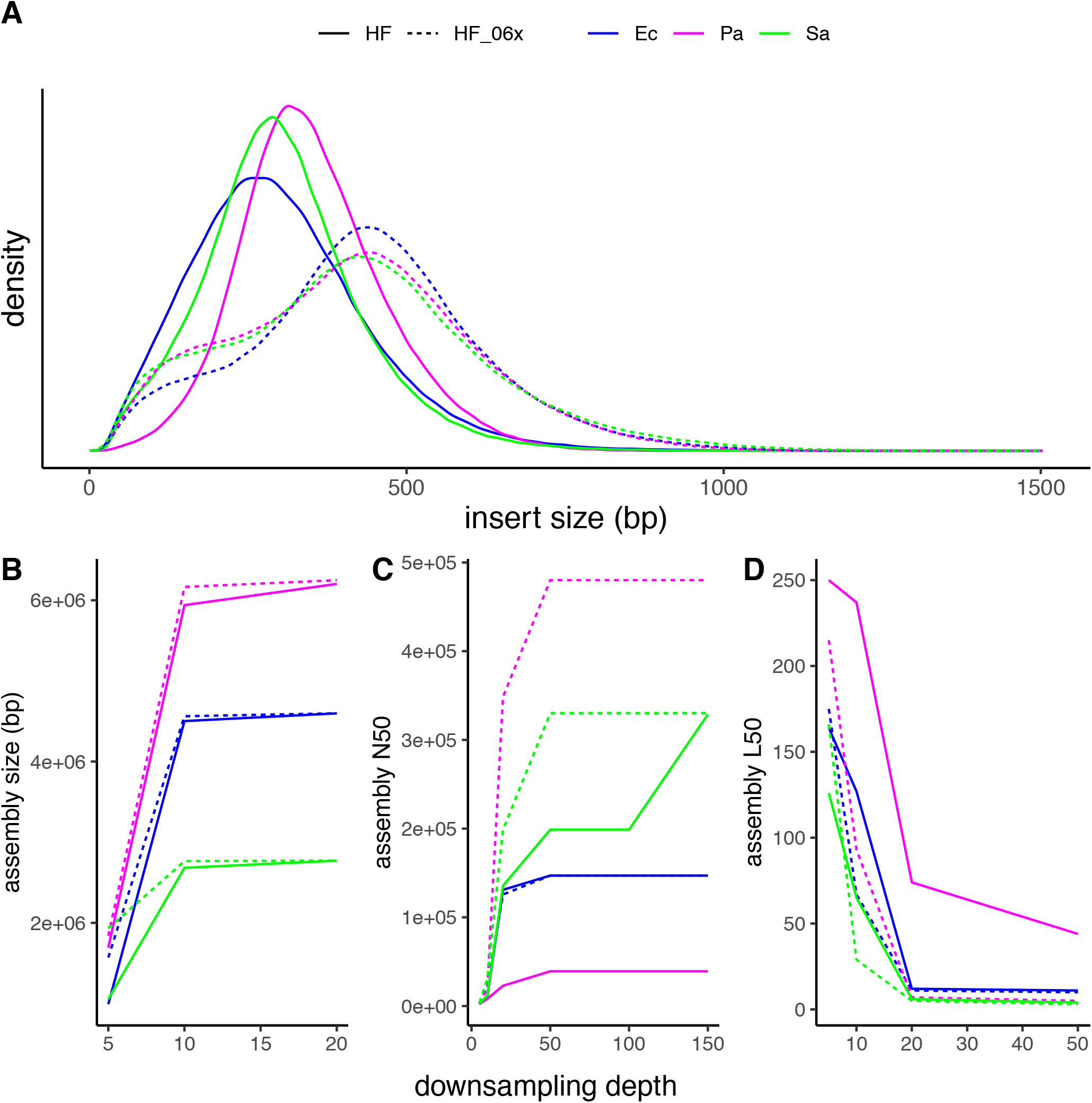
Insert size distribution and assembly metrics of double-left clean up Hackflex libraries compared to Hackflex libraries. Insert size distribution (**A**) are shown for Hackflex and double-left clean up Hackflex libraries from *E. coli* (blue) (Ec.HF.B3 median=287 mean=300.68; Ec.HF_06x.B3 median=433 mean=432.72), *P. aeruginosa* (pink) (Pa.HF.B2 median=341 mean=352.01; Pa.HF_06x.B3 median=417 mean=415.06), and *S. aureus* (blue) (Sa.HF.B2 median=296 mean=304.94; Sa.HF_06x.B3 median=416 mean=420.29) genomic DNA. The same libraries, down sampled to several depths, and assembled, were analyzed for assembly quality.:assembly size (**B**), and assembly metrics N50 (**C**), and L50 (**D**) are shown for Hackflex and double-left clean up Hackflex libraries from *E. coli* (blue), *P. aeruginosa* (pink), and *S. aureus* (blue).

To evaluate the effects of coverage depth as it relates to the influence of insert size, assemblies were performed from libraries downsampled to various coverage depths. Assemblies from Hackflex libraries were more fragmented (higher L50) compared to double left clean up Hackflex libraries. The effect was most prominent in libraries from *P. aeruginosa*. Resizing of the two libraries brings the depth to 33, and 34 mean coverage for Pa.HF_06x.B3 and Pa.HF.B2, respectively. Downsampling the two libraries to the common maximum depth achieved (33x), shows that a significant difference in assembly quality between protocols was detectable (Pa.HF.B2 N50=39,210 L50=44; Pa.HF_06x.B3 N50=480,098 L50=5). Similarly, the difference between the two protocols was prominent in libraries from *S. aureus*, where a larger N50 was obtained in the double left clean up Hackflex library compared to the Hackflex library at either 10x (Sa.HF.B2 N50=11,756 L50=65; Sa.HF_06x.B3 N50=30,473 L50=29), 20x (Sa.HF.B2 N50=135,652 L50=6; Sa.HF_06x.B3 N50=198,717 L50=5), or 50x (Sa.HF.B2 N50=198,879 L50=4; Sa.HF_06x.B3 N50=330,244 L50=3). Downsampling the two libraries to the common maximum depth achieved (median=116) shows that a similar assembly quality is reached (Sa.HF.B2 N50=328,703 L50=3; Sa.HF_06x.B3 N50=330,244 L50=3). In *E. coli* libraries, the benefit of larger insert sizes libraries on assembly quality was prominent at 10x depth (Ec.HF.B3 N50=10,798 L50=127 Ec.HF_06x.B3 N50=21,488 L50=67), mild at 20x depth (Ec.HF.B3 N50=131,011 L50=12 Ec.HF_06x.B3 N50=125,707 L50=11), and non-existent at 48x depth (Ec.HF.B3 N50=147,079 L50=11 Ec.HF_06x.B3 N50=147,079 L50=10). (**Figure 4**)

Other metrics were not affected by the double left clean-up. Equally re-sized libraries Ec.HF.B3 and Ec.HF_06x.B3 had a mean coverage of 46.9 and 46.8, respectively, with no sites with zero coverage. These libraries had a 0.66 correlation in coverage (*p*<0.0001), similarly to Ec.HF.B3 and Ec.SF_1:50.B2 (Pearson’s *R*=0.58; *p*<0.0001). Equally re-sized libraries Pa.HF.B2 and Pa.HF_06x.B3 had a mean coverage of 33.55 and 33.24, with 7,755 and 195 sites with zero coverage, respectively. These libraries had a 0.20 correlation (*p*<0.0001), similarly to Pa.HF.B2 and Pa.SF_1:50.B2 (Pearson’s *R*=0.14; *p*<0.0001). Equally re-sized libraries Sa.HF.B2 and Sa.HF_06x.B3 had a mean coverage of 117.03 and 116.09, respectively, with no sites with zero coverage. These libraries had a 0.42 correlation (*p*<0.0001), similarly to Sa.HF.B2 and Sa.SF_1:50.B2 (Pearson’s *R*=0.44; *p*<0.0001).

### Additional libraries

Additional libraries were prepared in order to test the effect of PrimeStar on the Standard Flex protocol, and the use of different annealing and extension temperatures in the Hackflex protocol. A lower annealing temperature was associated with a stronger GC bias in both *P. aeruginosa* (Pa.HF_55A.B2 *R*=-0.734 *p*<0.0001; Pa.HF.B2 *R*=-0.248 *p*=0.048) and in *S. aureus* libraries (Sa.HF_55A.B2 *R*=0.945 *p*<0.0001; Sa.HF.B2 *R*=0.741 *p*<0.0001). A lower annealing temperature and a higher extension temperature in the Hackflex protocol, only tested with *P. aeruginosa* genomic DNA, was associated with an even higher GC bias (Pa.HF_55A72E.B2 *R*=-0.851 *p*<0.0001; Pa.HF.B2 *R*=-0.248 *p*=0.048). (**Supplementary Table 2**)

## DISCUSSION

Our customised library preparation protocol, Hackflex, involves several modifications to the Nextera Flex method, including the use of a 1:50 dilution of the bead-linked transposase and the replacement of several kit components with alternative reagents to greatly expand the total number of libraries that can be produced from a single kit. The process of making the Hackflex reagents can take up to three hours. However, Hackflex reagents can be made in batches, with a batch used over a period of time of up to one year. To evaluate the performance of the Hackflex protocol, we constructed libraries in parallel using the standard Nextera Flex protocol (referred to as “Standard Flex”), an adapted version of the Nextera Flex protocol using the diluted transposase beads (referred to as “1:50 Flex”) and using the Hackflex protocol. Libraries were constructed from the genomic DNA of either *E. coli* MG1655, *P. aeruginosa* PAO1, or *S. aureus* ATCC25923.

### Barcode designs

To enable greater flexibility and higher multiplexing in sample barcoding, we designed our own barcode sequences, produced by IDT using their standard oligo plate manufacturing process. We generated two designs: v0 and v1. In the barcodes v0 design, i5 oligos are the tandem complement sequences of the i7 oligos. Although the tandem design of v0 barcodes did not significantly impact the quality of the data on a MiSeq instrument, a significant degradation of index read 2 quality (i5) was observed when such barcode combinations were applied to generate data for a large scale porcine gut metagenomics project^17,18^. It was noted by Illumina technical support that the tandem complement design could lead to significantly reduced quality scores for the i5 index read on the NovaSeq instrument (personal communication, Illumina). We therefore designed new barcodes, which we refer to as barcodes v1, that do not have a tandem complement design and we tested their performance by constructing 96 Hackflex libraries, from 96 barcode combinations.

Incorrect sample barcode assignment can occur either due to barcode cross-contamination during sample preparation or sequencing, or it can be due to errors during sequencing such as the presence of multiple overlapping clusters on the flow cell surface, base miscalls, and image processing errors during cluster calling. Sample barcode misassignments can be minimised if sample barcode oligos are designed to be sufficiently unique^19^, where more than 3 bp must be errors for the barcode to be misallocated. The use of unique dual indexing where each of i5 and i7 index sequences are unrelated to each other is also known to mitigate sample barcode misassignment issues (https://support.illumina.com/bulletins/2018/08/understanding-unique-dual-indexes--udi--and-associated-library-p.html). From our v1 barcodes, we report a barcode cross-contamination rate of 0.002 for our new barcodes. How this translates into sample misassignment of reads will depend heavily on the number of samples in a pool and the specific barcode oligos in use. This low rate of sample misassignment should be sufficient for consensus sequencing applications such as de novo genome assembly, where the impact of a low rate of misassigned reads is likely to be very limited.

Three out of the ninety-six v1 barcode libraries yielded a low read count. One possible cause is pipetting error during various stages of the library prep where a multichannel pipette was used. Alternatively, failures could be attributable to barcode-specific reaction kinetics. One of these libraries with low read counts has an i7 barcode for which a particularly low deltaG is predicted for homodimer formation, but the remaining i5 and i7 barcodes appear indistinguishable from other barcodes in terms of GC content, hairpin, homodimer, and heterodimer thermodynamics.

While a significant negative correlation was found in the v0 design, no correlation was found in the v1 barcode design between the barcode GC content and the read count obtained. There are several possible explanations for the variance in barcode representation in our design, including formation of homo- and heterodimers, and hairpins. The v1 barcodes have been designed to be distinct from the current set of standard 8bp Illumina barcode sequences. There is no reason a user of the Hackflex protocol would not be able to use standard Illumina barcode adapter oligos together with our protocol modifications, however we have not tested performance under those conditions.

Worthy of note, the performance of the Hackflex barcode libraries is assessed based on the read count obtained from each barcode set, which can be affected by the barcode oligo performance as well as by the lab operator. We believe that the observed performance can be interpreted as a lower bound on the performance of the oligos themselves. That is, in the absence of operator error, the numbers reflect the performance of the oligos, but the true oligo performance could be better given that some of the observed variation may have been introduced by the operator.

### Quality of sequence data

Hackflex uses a lower amount of input DNA (10 rather than 200 ng) compared to the standard Flex protocol. Coupled to the reduction of input DNA, a higher number of PCR cycles is used in Hackflex (12 instead of 5). This modification could be expected to increase the number of PCR duplicates. However, we did not detect a higher number of PCR duplicates in diluted libraries (*i.e*.: Hackflex and 1:50 Flex) compared to undiluted libraries (*i.e*.: standard Flex), suggesting that diluted libraries are either equally efficient as undiluted libraries, or that any apparently duplicate read pairs are derived from transposition bias or flowcell duplicates. It is likely that much deeper sequencing of these libraries would be required to reveal differences in library complexity. Raw libraries from distinct protocols (*i.e*.: standard Flex, 1:50 Flex, and Hackflex) showed comparable quality scores and behaviour through the adapter and quality trimming steps. To further compare the performance of distinct protocols, libraries were randomly subsampled to achieve equally sized libraries and undergo mapping against a reference genome.

As a consequence of the polymerase’s bias against regions with extreme GC content, it can be expected that by increasing the PCR cycles, GC bias will be increased. The GC bias was lower for Hackflex libraries than for Standard Flex and/or 1:50 Flex, in *S. aureus* and *P. aeruginosa*, respectively. For *E. coli*, we were able to provide an estimate of GC bias from the top 20 Hackflex barcode v1 libraries with the highest coverage depth. The GC bias estimate obtained from these libraries had a mean GC bias of −0.642 (sd=0.043), which is lower than that of standard Flex and diluted Flex libraries from batch 1, and higher than that of standard Flex and diluted Flex libraries from batch 2. However, as the GC bias estimates for standard Flex and 1:50 Flex from *E. coli* genomic DNA were derived from 4 libraries (2 replicates), and the number of libraries for direct comparison for *P. aeruginosa* and *S. aureus* is also limited, so we could not draw a statistically significant conclusion that Hackflex has a lower or equally low GC bias compared to standard Flex.

Additionally, we explored other conditions for Hackflex, such as a lower annealing and a higher extension temperature. This resulted in a stronger GC bias but the limited number of datapoints means the result is inconclusive. An effect on the uniformity of coverage could have been expected when substituting PrimeStar for EPM in the Hackflex protocol. Although a batch effect was detected, the coverage obtained with the distinct protocols was highly similar within each batch, as indicated by the read mapping rates, the number of unmapped reads, the median and average coverages, and the number of sites reporting zero coverage. Sites with zero coverage in *E. coli* and *S. aureus* were located near the extremities of contigs, raising the possibility that the coverage dropouts may be due to read mapping artefacts rather than lack of sequence data. However, a higher number of sites with zero coverage (double) were obtained for *P. aeruginosa* with Hackflex, as compared to the standard Flex protocol. We are unsure how to explain this observation given that PrimeSTAR GXL is specified for PCR of high GC samples such as *P. aeruginosa*. However, the fact that the reference genome (against which the libraries were mapped) was assembled from the standard Flex and diluted Flex libraries could have biased the mapping in favor of those libraries and against Hackflex. Other coverage metrics were highly similar among the different protocols, and Hackflex showed nearly a fourfold lower GC bias on the high GC *P. aeruginosa* genome than either standard Flex or 1:50 Flex.

We analysed insert size distributions obtained with Hackflex and compared it to that obtained with standard Flex and 1:50 Flex. We did not find significant differences in insert size distributions between distinct protocols. However, as insert size distribution is highly dependent on manual pipetting, and could also be affected by undocumented BLT chemistry changes across distinct batches, it is difficult to draw any conclusions from comparisons of insert size distributions between protocols across batches.

### Tunability to other size ranges

The size distribution of fragments generated by the standard Nextera Flex workflow appears to be strongly skewed towards small fragments. This is seen in Hackflex as well, and is likely driven by the current formulation of bead-linked transposase. While the fragment sizes generated by tagmentation with transposase in-solution are highly sensitive to the relative concentrations of DNA and transposomes, the fragment sizes generated from BLT appear much less sensitive to those parameters. This has the benefit of making the reaction more robust to variation in input DNA concentration, but comes at a cost in that there is less flexibility to optimize the yield of fragments in a particular size range. In particular, when carrying out *de novo* assembly, longer reads from larger inserts are known to have a beneficial effect^20^. Moreover, longer reads such as the PE250 mode offered on MiSeq and Novaseq instruments may offer less benefit on inserts less than 500nt in size, because PE250 on shorter inserts would generate overlapping reads on the same fragment, yielding redundant sequence data. Therefore, we evaluated the ability of Hackflex to produce larger insert libraries of sufficient complexity for *de novo* microbial genome assembly.

Hackflex libraries were made from *E. coli, P. aeruginosa* and *S. aureus*, where two rounds of bead cleanup were carried out to remove the small insert size. This is done by cleaning up the libraries through SPRI beads size selection at 0.6x ratio twice (double-left clean up). In doing so, the fragments with small insert were removed and the average insert size length of the library has increased. The increase of insert size can be of benefit for assembly, and particularly, for assembly of certain species. As we ran assemblies at different depths, we compared the assemblies obtained, and concluded that the double left clean up step in Hackflex helps improve the assembly, in particular at lower coverage and for *P. aeruginosa*, which is a larger and more complex genome than the other two. Despite improving the assembly, the double left clean up did not affect the quality of the data produced, as reported by other metrics.

### Applicability to large and complex genomes

Given that the transposase dilution has a direct effect on library complexity, the Hackflex method may not be suitable for very large genomes such as those of humans and plants. In order to increase the library complexity the Hackflex method could be tweaked to decrease the transposase dilution. However, Hackflex is designed to enable large scale sequencing projects for low complexity genomes such as bacteria, including metagenomic projects with high sample numbers^17,18^ or DNA surveillance projects, where thousands of bacterial genomes must be sequenced.

## CONCLUSION

Here we have developed and characterised an alternative method of library construction for Illumina sequencing which by reducing the library prep expenses, allows users to process from 9.87-fold to 14-fold more samples at the same reagent cost. This study demonstrates that data of comparably high quality can be obtained with Hackflex as could be generated by the existing Nextera Flex method. Comparison with the existing Nextera Flex method demonstrates that Hackflex is a valid and cost-effective alternative to construct libraries at a large scale.

## Author contributions

Conceived and designed the experiments: DG KA ML JT LM AD. Performed the experiments: DG KA ML JT LM. Analyzed the data: DG AD. Contributed reagents/materials/analysis tools: KA JT ML LM. Wrote the manuscript: DG. Edited the manuscript: AD KA LM.

